# Prevalence of coronary artery disease risk factors among fire-fighters in Cape Town, South Africa

**DOI:** 10.1101/652503

**Authors:** Ghaleelullah Achmat, Rucia V. November, Lloyd L. Leach

**Affiliations:** Department of Sport, Recreation and Exercise Science, University of the Western Cape, Cape Town, Western Cape, South Africa

## Abstract

**Background:** Cardiovascular disease is a major cause of morbidity and on-duty mortality among fire-fighters. This study investigated the prevalence of coronary artery disease (CAD) risk factors among firefighters in Cape Town, South Africa.

**Methods:** A quantitative, cross-sectional and descriptive study design was used. A convenient sample of 219 male fire-fighters with mean age 37.85±9.80 years was recruited. Eight major CAD risk factors were measured. SPSS (ver. 23) was used with the Pearson correlation and Kruskall-Wallis H test with the Mann Whitney test post hoc and a Bonferroni correction.

**Results:** The significance level set at p<0.05. Most fire-fighters (65.29%) were stratified as moderate risk for CAD, with 21.00% as low risk, and 14.15% as high risk. A sedentary lifestyle was the most prevalent CAD risk factor (51.14%), followed by obesity (45.90%). and cigarette smoking (38.30%). Statistically significant correlations were found between waist circumference and body mass index (BMI) (r=0.711; p<0.01), hip circumference and BMI (r=0.673; p<0.01), waist circumference and waist-hip ratio (WHR) (r=0.665; p<0.01), and BMI and body mass (r=0.512; p<0.01).

**Conclusion:** The majority of fire-fighters presented with multiple modifiable CAD risk factors. Fire departments should focus on promoting education on CAD to reduce the risk of fire-fighters developing heart disease.

## Introduction

Fire-fighters need good cardiorespiratory health and fitness to be able to respond to the job’s demanding emergency situations [1]. Globally, it is understood that fire-fighting is a physically challenging occupation [2]. Studies have shown that coronary artery disease (CAD) is more prevalent among fire-fighters than any other occupation in the United States [3]. Approximately 50 percent of fire-fighter line-of-duty-deaths (LODDs) occur as a result of heart attacks [3], [4]. These cardiovascular events do not only happen at random, but mostly occur during very physically demanding situations, such as fire suppression [4].

Furthermore, the majority of fire-fighters who experienced these events possess one or more of the following risk factors for CAD, namely, obesity, prediabetes, dyslipidemia, high blood pressure (hypertension), cigarette smoking, and/or a sedentary lifestyle. The risk of heart attacks among fire-fighters is dependent on many factors, including factors such as a low level of physical activity, and chronic exposure to smoke. In addition, the high temperatures present during fire suppression significantly increase physiological strain and the potential to over exert the cardiovascular system [4].

Many countries in Africa bear a heavy burden from cardiovascular disease, more especially in sub-Saharan Africa [5]. In South Africa, the prevalence of cardiovascular disease has been aggravated by an increased burden of cardiovascular risk factors, such as cigarette smoking, hypertension, dyslipidemia, diabetes and sedentary lifestyles [6]. The majority of individuals affected are young and at their most productive age, and constitute the largest sector of the workforce, fire-fighters included [6].

Many of the fire-fighters, who died on-duty, did not have an up-to-date medical evaluation [2]. Hypertension, prior occlusive disease, and cigarette smoking presented as significant risk factors for on-duty-death [2]. Furthermore, some of the firefighters who were older than 45 years had up to six times the risk of cardiovascular disease [7]. Due to the scarcity of research on CAD risk factors among fire-fighters, especially in Africa, the purpose of this study was to determine the prevalence of CAD risk factors among fire-fighters in the Western Cape Province, South Africa.

## Methods

### Research Design

A quantitative cross-sectional and descriptive design was used in this study.

### Recruitment of Participants

A convenient sample of 219 male career fire-fighters aged 18–65 years from the Western Cape Province, South Africa, gave their written informed consent to participate in the study. Participants were excluded if they were administrative staff and volunteer fire-fighters (i.e., fire-fighters contracted for the peak fire season only).

### Data Collection

Each fire-fighter completed a researcher-generated CAD risk factor questionnaire. Risk stratification was determined according to the guidelines of the [8].

Physical measurements included stature and body mass using a combination stadiometer and beam balance scale (Seca model 700, Gmbh & Co., Germany). Hip and waist circumferences were measured using a non-distensible metal tape measure (Sanny Medical,rk HK). All physical measurements were done according to the guidelines of the International Society for the Advancement of Kinanthropometry (ISAK) [9]. Blood pressure was measured with a pressure cuff of appropriate size (12 cm × 35 cm) using a standard mercury sphygmomanometer (Goodpro International Co., Limited, China) and an acoustic Sprague Rappaport stethoscope (Medical Supplies and Equipment Company, Houston, Texas, USA). Total cholesterol was measured with an Accutrend Plus meter (Roche Diagnostics, 04235643059, GmbH, Germany, and fasting blood sugar with an AccuCheck meter (Roche Diagnostics, 95346754160, GmbH, Germany) using a micro-capillary blood sample from a finger-prick. The researchers calibrated all research instruments.

### Data Analysis

The Statistical Package for the Social Sciences (SPSS), version 23 (IBM, New York, USA) was used for data analysis. Descriptive statistics (mean and standard deviation) and inferential statistics (Pearson product-moment correlation coefficient, and the Kruskal Wallis H test was applied between groups with the Mann Whitney test post hoc and a Bonferroni correction was applied to the post hoc analysis to minimise type I error, so that all effects are reported at a 0.0167 level of significance [10]. A p<0.05 indicated statistical significance.

### Ethical Considerations

The Senate Biomedical Research and Ethics Committee (BMREC) of the University of the Western Cape granted ethical clearance to conduct the study (Ethics clearance number: 14/10/52). Permission was also granted by the Chief Fire Commander of the City of Cape Town Fire and Rescue Service. An information letter was issued to all fire-fighters and their consent to participate voluntarily was requested in writing.

## RESULTS AND DISCUSSION

The results of the physical measurements and CAD risk factors are presented according to different weight categories in Table 1 as mean (±SD). Body mass was significantly different between the three groups [H(2)=55.705, p=0.000]. Post hoc analysis displayed significant differences between normal weight and obese groups (U=1597, r=−0.53), and overweight and obese groups (U=908, r =−0.38).

**Table 1: Physical characteristics of fire-fighters based on three weight categories.**

Body mass index (BMI) differed significantly between groups [H(2)=187.28, p=0.00]. Post-hoc analysis displayed significant differences between normal and overweight groups (U=0.00, r=–0.79), normal and obese groups (U=0.00, r=-0.87), and overweight and obese groups (U=0.00, r=–0.79).

Waist circumference was significantly different between groups [H(2)=89.870, p=0.000]. Post hoc analysis showed significant differences between normal weight and overweight groups (U=850.5, r=–0.39), normal weight and obese groups (U=882.5, r=−0.68), and overweight and obese groups (U=933, r = −037).

Hip circumference was significantly different between groups [H(2)=93.639, p=0.000]. Post hoc analysis displayed significant differences between normal weight and overweight groups (U=933, r=–0.35), normal weight and obese groups (U=808, r=−0.69) and overweight and obese group (U=844, r=−0.41).

Waist-hip ratio was significant different between groups [H(2)=17.530, p=0.000]. Post hoc analysis displayed significant differences between normal weight and overweight groups (U=1210, r=–0.22), and normal weight and obese groups (U=2679, r=−0.29).

Systolic blood pressure was significantly different between groups [H(2) = 25.159, p = 0.000]. Post-hoc analysis showed significant differences between normal weight and obese groups (U=2435, r=–0.35), and overweight and obese group (U=1123.5, r=−0.29).

Diastolic blood pressure was significantly different between groups [H(2)=21.484, p=0.000]. Post hoc analysis displayed significant differences between normal weight and obese group (U=2513.5, r=–0.33.

Fasting blood glucose (prediabetes) was significantly different between groups [H(2)=14.136, p=0.001]. Post hoc analysis displayed significant differences between normal weight and overweight groups (U=1211.5, r=–0.26), and normal weight and obese group (U=3160, r=-0.26).

Waist circumference (WC) had a significant relationship with BMI (r=0.711, p<0.01), waist-hip ratio (r=0.665, p<0.01), systolic blood pressure (SBP) (r=0.414, p<0.01), diastolic blood pressure (DBP) (r=0.396, p<0.01), and body mass (BM) (r=0.389, p<0.01). Waist-hip ratio (WHR) had a significant relationship with hip circumference (HC) (r=0.665, p<0.01), SBP (r=0.414, p<0.01), and DBP (r=0.377, p<0.01). BMI had a statistically significant relationship with HC (r=0.673, p<0.01), BM (r=0.512, p<0.01), SBP (r=0.330, p<0.01), DBP (r=0.313, p<0.01), and WHR (r=0.313, p<0.01). Body mass had a significant correlation with HC (r=0.385, p<0.01).

### Prediabetes

Fig. 1 showed that 17.27% of fire-fighters presented with prediabetes. Prediabetes and obesity affected work performance of fire-fighters work making it difficult to climb ladders, especially when loaded with equipment [11]. Among 5065 fire-fighters screened for fasting glucose, 5.9% were prediabetic and 2.7% diabetic [12]. In a cohort study of 957 career fire-fighters, approximately 26% displayed high blood glucose levels [13]. Similarly, in 2010, among 216 fire-fighters medically screened, 32.9% were prediabetic and 14.8% were diabetic [14]. Obesity coupled with physical inactivity is related to the risk of developing diabetes and CVD [12]. The prevalence of type II diabetes among fire-fighters is linked with a 21% risk of experiencing an on-duty CAD event [4].

**Fig. 1.**
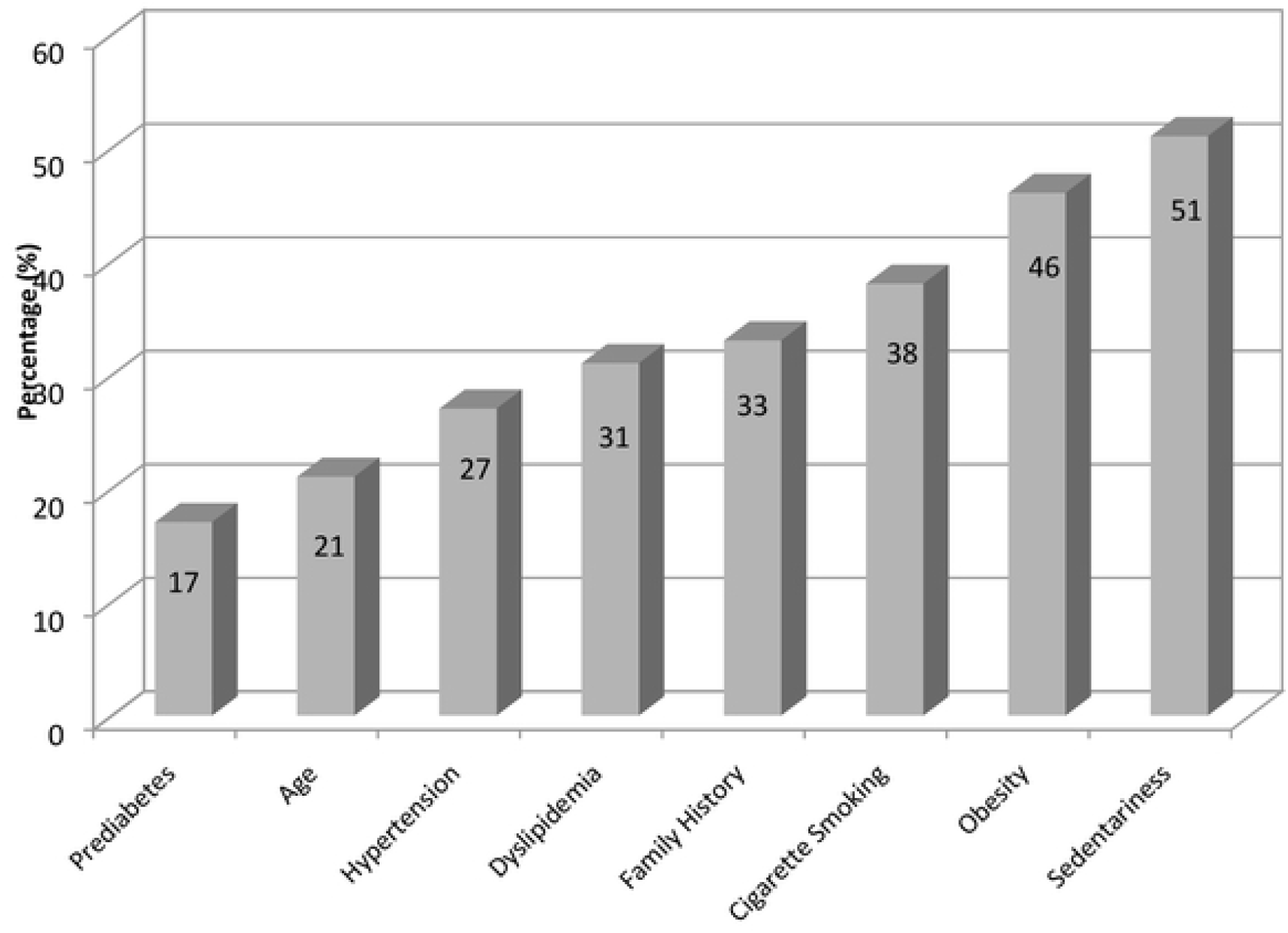
Prevalence of CAD risk factors among fire-fighters.

### Age

Age is associated with increased risk of CAD in men aged 45 years and older (ACSM, 2014, p. 28). In Fig. 1, 20.51% of fire-fighters were above the age of 45 years that placed them at moderate risk for CAD. Furthermore, physical activity tends to decrease with age and is less prevalent amongst those with chronic disease [11].

### Hypertension

Hypertension is common in the South African population [6]. Hypertension was found in 27.30% of fire-fighters, and 9.41% were pre-hypertensive (Fig. 1). In the present study, blood pressure correlated significantly with body composition, i.e., WC, WHR and BMI, which together exponentially increased the risk of CAD [1], [11]. For every 20 mm Hg increase in systolic blood pressure or 10 mm Hg increase in diastolic blood pressure, the risk of CVD increases twofold [15]. Alarms, sirens, vehicle engines, and mechanized rescue equipment typically produce average noise exposures in the 63 to 85 decibel (dBA) range that increase blood pressure [3]. Siren noises elevate systolic blood pressure by 5.9 to 11.8 mm Hg [3].

Hypertension is an independent predictor of adverse outcomes, such as injury on duty, termination of duty, resignation, premature retirement and cardiovascular events, including on-duty death [11]. Chronically uncontrolled hypertension and hypertensive heart disease were responsible for nearly 8% of cardiovascular disability retirements [11], [16].

### Dyslipidemia

Dyslipidemia was prevalent among 31% of fire-fighters, and has been associated with excess adipose tissue, glucose intolerance, hypertension and endothelial dysfunction, all of which affect the hearts function [1], [12], [15]. In a longitudinal study from 2004–2007, 37% of 7904 fire-fighters had dyslipidemia [14]. Like most shift workers, fire-fighters are more likely to eat fast-foods or ‘quick’ meals with high proportions of fat that adversely affect blood lipid profiles and accelerate the atherosclerotic process [17].

### A Family History of Heart Disease

A family history of CAD was present in 32.80% of fire-fighters. The risk of CAD is exponentially increased when family history is combined with poor dietary habits, physical inactivity and smoking [18]. Family history also plays a major role in the development of prediabetes and type 2 diabetes [16], [18].

### Cigarette Smoking

Between 1987–1994, the average prevalence of cigarette smoking amongst fire-fighters was 26.9% [19]. In the present study, cigarette smoking was prevalent in 38.30% of the fire-fighters, and is associated with other significant health and safety risks [20]. Exposure of fire-fighters to carbon monoxide contributed to the increased risk of heart disease and stroke [21]. Smokers were more likely to be diagnosed with anxiety disorder, alcohol problems and found driving under the influence than non-smokers [20].

### Obesity

Being obese is a national epidemic in South Africa and a major risk factor for CAD [6]. Fire-fighters experience high rates of obesity ranging from 30%-40%, similar to that found among the general adult population (Flegal, 2010). The current study showed that 45.90% of fire-fighters were obese with high waist circumference measurements that predisposed them to increased cardiovascular events [1], [3], [4]. Overweight and obese fire-fighters often presented with multiple CAD risk factors, such as prediabetes, hypertension, and hypercholesterolemia [1], [11], [16], [17], [22]. Fire-fighters with a high BMI were at increased risk for work-related injuries and disabilities [4].

Fire-fighters experience weight gains of approximately 0.5 to 1.6 kg per year throughout their career [23]. Furthermore, the risk for diabetes increased by 4.0% for every centimetre gained around the waist [24]. Abdominal obesity, a sedentary lifestyle, hormonal imbalances, insulin resistance, and a poor diet are the prime factors in the development of metabolic syndrome [21].

### A Sedentary Lifestyle

In the present study, a sedentary lifestyle was the most prevalent CAD risk factor, occurring in 51.14% of fire-fighters. This places fire-fighters at risk, because they do not meet the minimum physical activity requirements for health benefits, i.e., participating in at least 30 minutes of moderate-intensity (40%-60% VO2max) physical activity most days of the week [8].

Physical activity is very important to fire-fighters as a public service, with work demands placing significant stress on the cardiovascular system [4]. During intense work, fire-fighters heart rates were reported to reach between 70 and 90% of their age-predicted maximal heart rates [25]. This, coupled with the sudden change from long periods of relative inactivity to sudden high-intensity work, while wearing relatively weighted protective fireproof clothing, increases the risk of morbidity and mortality in fire-fighters [26]. High levels of physical fitness enable fire-fighters to perform their job safely or effectively [4], and mitigate the impact of other cardiac risk factors, such as obesity and diabetes [21]. The irregular bouts of prolonged vigorous and strenuous activity required in fighting fires are a precursor to acute coronary heart events, especially in fire-fighters with low fitness levels [4].

### Relationship Between Various CAD Risk Factors

In Fig. 2, 9% of fire-fighters had zero risk factors, whereas 12% had one risk factor only, and are stratified as low CAD risk. Moderate CAD risk stratification includes two or more risk factors. Fire-fighters presenting with 2, 3, 4, 5, 6, 7, and 8 risk factors were 25%, 22%, 16%, 7%, 6%, 2% and 1%, respectively. Most importantly, the presence of advanced atherosclerotic lesions and pathological changes are directly correlated with the number of risk factors [3], [4], [13], [14].

**Fig. 2.**
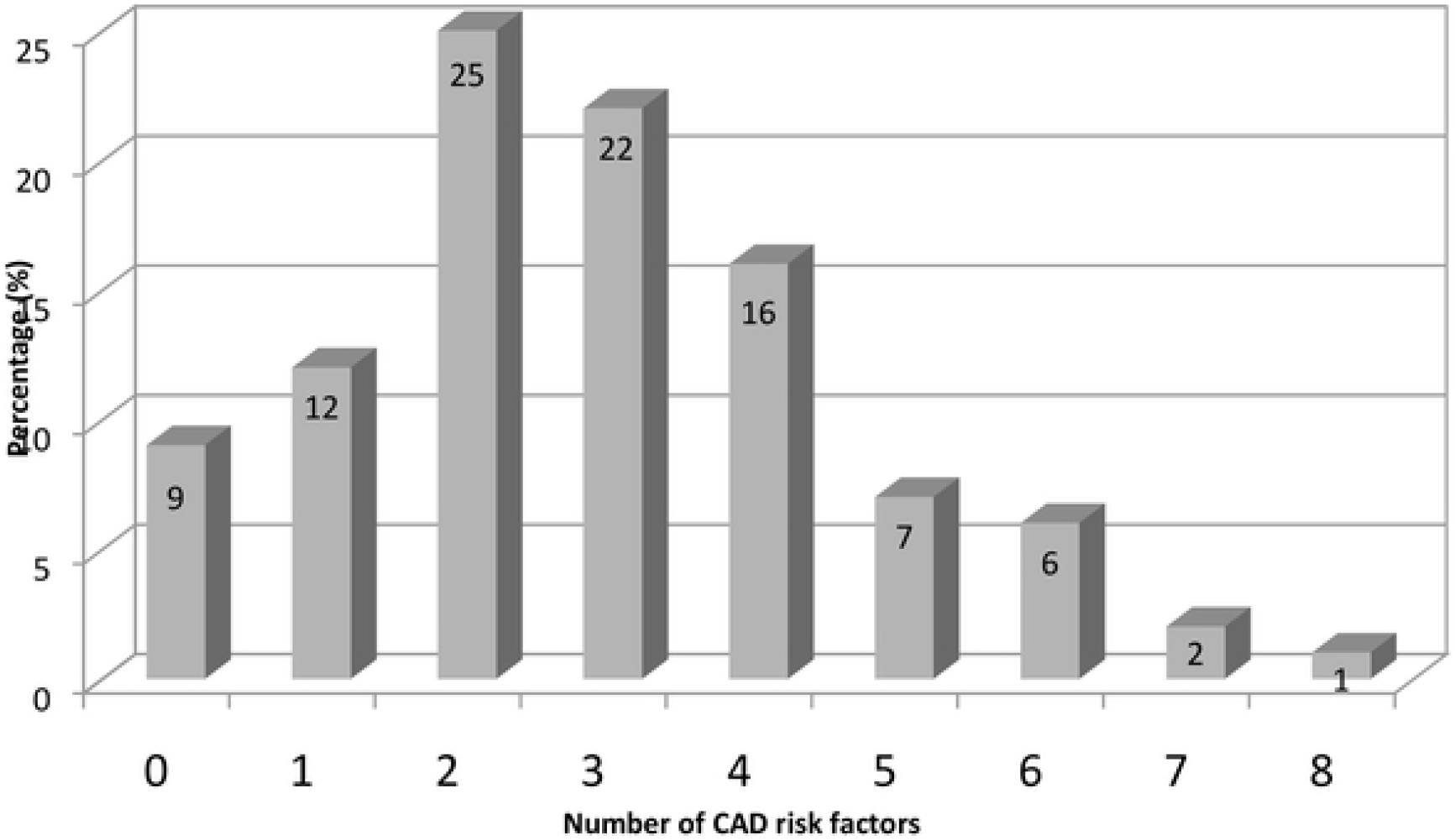
Frequency of CAD risk factors among fire-fighters.

Understanding the major independent CAD risk factors is critical for the fire service, due to the impact of CAD on job performance and on-duty deaths [3], [4]. Risk factors are the result of interactions between genetic and environmental factors over an extended period of time [27].

### CAD Risk Stratification

Fig. 3 shows that 64.4% of fire-fighters were at moderate risk for CAD, 21.0% were at low risk, and 14.6% were at high risk, particularly the older fire-fighters. In the 45 male fire-fighters aged 45 years and older, 53.33% were at high risk (i.e., diagnosed with chronic disease). An increase in age is an independent predictor of CAD in fire-fighters [4].

**Fig. 3.**
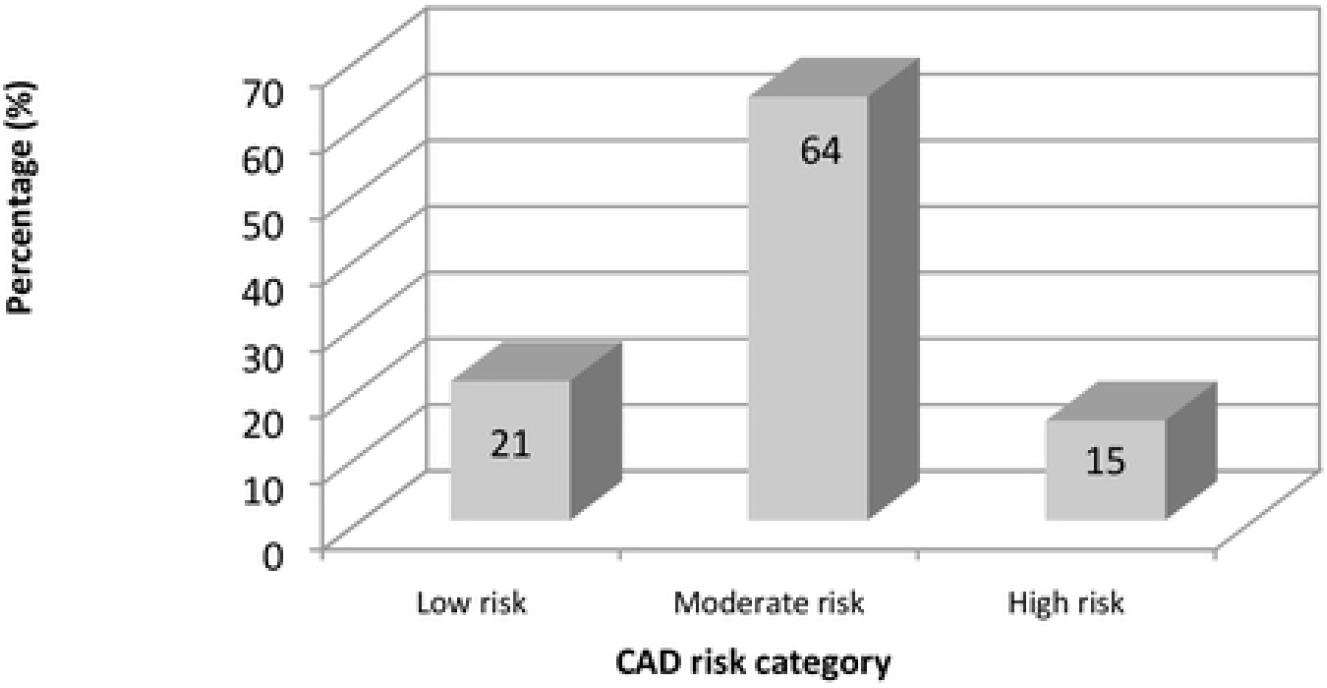
Coronary artery disease (CAD) risk stratification of fire-fighters.

### Strengths of the Study

There is very little research done on coronary artery disease risk factors in fire-fighters globally and, more especially, very little throughout Africa, and South Africa specifically. The study highlights the burden of coronary artery disease among firefighters in the City of Cape Town in terms of fitness for duty and public safety. It is an important public health and safety issue in the context of Africa and South Africa, and this study adds to the limited research done on this topic.

### Limitations of the Study

The study did not use randomized sampling that impacted the external validity of the findings. The study did not include female fire-fighters and, therefore, has limited application with regard to female fire-fighters.

## Conclusion

The majority of fire-fighters screened in this study were stratified as moderate risk for CAD, due to the presence of multiple modifiable (preventable) risk factors such as sedentariness, obesity and cigarette smoking. A sedentary lifestyle was the most prevalent risk factor amongst fire-fighters, which is cause for concern from a personal and public health perspective. There is strong consensus that fire-fighting is a physically demanding and stressful occupation that requires a good level of cardiovascular fitness [4]. The frequency of cardiac events amongst firefighters is reported often in the literature [3], [4], [14], [16], and is cause for concern, especially since the risk factors are well-known, detectable, and largely preventable. Early risk assessment and disease prevention efforts are lacking in this occupational group, particularly in South Africa [6].

## Acknowledgments

The City of Cape Town Fire and Rescue Service is acknowledged for their willing participation in the study.

## Supporting information

**S1 Fig. This is the S1 Fig 1 Title. Prevalence of CAD risk factors among fire-fighters.**

**S2 Fig. This is the S2 Fig 2 Title. Frequency of CAD risk factors among fire-fighters.**

**S2 Fig. This is the S2 Fig 3 Title. Coronary artery disease (CAD) risk stratification of fire-fighters.**

**S1 Table. This is the S1 Table 1 Title. Physical characteristics of fire-fighters based on three weight categories**

